# Cholinergic upregulation by optogenetic stimulation of nucleus basalis after photothrombotic stroke in forelimb somatosensory cortex improves endpoint and motor but not sensory control of skilled reaching in mice

**DOI:** 10.1101/2020.06.09.143354

**Authors:** Behroo Mirza Agha, Roya Akbary, Arashk Ghasroddashti, Mojtaba Nazari-Ahangarkolaee, Ian Q. Whishaw, Majid H. Mohajerani

**Affiliations:** Department of Neuroscience, Canadian Centre for Behavioural Neuroscience, University of Lethbridge, Lethbridge, Alberta, Canada, T1K 3M4; Department of Psychology, University of Toronto, Toronto, Ontario, Canada, M5S 1A1

**Keywords:** acetylcholine upregulation, optogenetic stimulation of the nucleus basalis cholinergic neurons, photothrombotic stroke, sensory neglect, skilled reaching task in mice

## Abstract

A network of cholinergic neurons in the basal forebrain innerve the forebrain and are proposed to contribute to a variety of functions including attention, and cortical plasticity. This study examined the contribution of the nucleus basalis cholinergic projection to the sensorimotor cortex on recovery on a skilled reach-to-eat task following photothrombotic stroke in the forelimb region of the somatosensory cortex. Mice were trained to perform a single-pellet skilled reaching task and their pre and poststroke performance, from Day 4 to Day 28 poststroke, was assessed frame-by-frame by video analysis with end point, movement and sensorimotor integration measures. Somatosensory forelimb lesions produced impairments in endpoint and movement component measures of reaching and increased the incidence of fictive eating, a sensory impairment in mistaking a missed reach for a successful reach. Upregulated acetylcholine (ACh) release, as measured by local field potential recording, elicited via optogenetic stimulation of the nucleus basalis improved recovery of reaching and improved movement scores but did not affect a sensorimotor integration impairment poststroke. The results show that the mouse cortical forelimb somatosensory region contributes to forelimb motor behavior and suggest that ACh upregulation could serve as an adjunct to behavioral therapy for the acute treatment of stroke.

## Introduction

Stroke is a major cause of death and disability that results from a transient or permanent reduction in cerebral blood flow ^1^. During stroke, neuronal structure and function are rapidly damaged ^2^, but can in part recover during reperfusion ^3^ or following treatment agents that interrupt the events leading to cell death ^4, 5^ or induce hypothermia ^6^. Although neuronal structure can recover after intervention, balance of excitation and inhibition is altered ^7, 8^ and brain activity is depressed ^9^ which actively impede poststroke functional reorganization ^10-12^. The cellular and biochemical changes related to recovery following cortical stroke include changes in ionic balance, alterations in the properties of the cell membrane, changes in second messenger and mRNA production resulting in protein production and inflammatory response, and alterations in neurotransmitter function ^13-16^. Included in the alterations in neurotransmitter function, research suggests that acetylcholine (ACh) innervation in the neocortex may be important for recovery because evidence suggests that ACh is related to cortical plasticity ^17-23^, including the plastic changes that mediate recovery/compensation in the acute period after cerebral stroke ^24^. Accordingly, the acute poststroke period has been the target of a number of attempts to enhance behavioral recovery in the mouse using treatments that directly or indirectly enhance ACh activity ^25-28^. Despite the success of these treatments, the contribution and specificity of ACh to recovery has not been clarified, but one explanation is that ascending basal forebrain cholinergic system is involved ^29^. Previous findings show that, direct and indirect activation of excitatory neurons within peri-infarct circuits using optogenetic stimulation can promote functional recovery ^30, 31^. To further study the role of ACh in recovery/plasticity following cortical stroke, the cholinergic nucleus basalis (NB) neurons, the source of cholinergic projections to the sensorimotor region of the neocortex ^32, 33^ were ontogenetically stimulated daily during a portion of the acute recovery period (poststroke day 4 till day 14). Stroke in the forelimb region of the somatosensory cortex was induced by photothrombosis, a noninvasive method to produce a focal infarct that is consistent in terms of size and location. Because an acute recovery from stroke has been a target of study in experiments of functional recovery of mice performing skilled reaching tasks, the present study examined performance during poststroke period in mice trained on the single pellet reaching task. Endpoint measures of success, movement component scores, and sensorimotor integration were obtained for daily reaching tests. The analyses indicate that optogenetic stimulation of the nucleus basalis cholinergic neurons during the poststroke period improved some measures of recovery on end point measures and movement but was without effect on measures of sensory integration

## Material and Methods

### Animals

Twenty-seven adult ChAT-Cre/Ai32(ChR2-YFP), referred to hereafter as ChAT-Cre/Ai3 (16 male, 11 female) mice (3-4 months of age), weighing 20-30 g, raised at the Canadian Centre for Behavioral Neuroscience vivarium at the University of Lethbridge, were used. The ChAT-Cre/Ai32 line was generated by crossing ChAT-Cre (B6;129S6-Chat^tm1(cre)Lowl^/J; Jax stock number 006410) mouse line with Cre-dependent reporter lines Ai32 (129S-Gt(ROSA)26Sor^tm32(CAG-COP4^*^H134R/EYFP)Hze^/J; Jax stock number 012569) ^34^. This crossing is expected to produce expression of ChR2 in cholinergic neurons ^35^. The animals were housed in littermate quads after weaning on a 12h:12h light/dark cycle with light starting at 7:30am and temperature set at 22 °C. After fiber optic and LFP electrode implantation, the mice were housed singly. Testing and training were performed during the light phase of the cycle in the morning each day. Procedures were approved by the University of Lethbridge Animal Care Committee in accordance with the guidelines of Canadian Council on Animal Care. Mice were randomly assigned in three groups: stroke + stimulation (n = 9; 5M, 4F), stroke + no stimulation (n = 8; 5M, 3F), and sham stroke + no stimulation (n = 10; 6M, 4F). All the mice underwent identical handling and behaviour and stimulation procedures such that even animals in the No Stim and Sham Groups were brought to the stimulation room and were attached to the laser cable for the same duration as the Stim Group but did not receive laser stimulation.

### Animal surgery

To implant the fiber optic and LFP electrode (Figure 1A), animals were anesthetized with isoflurane (1-2%), and stereotaxic surgery was conducted using aseptic methods. The mice were placed in a stereotactic frame (Kopf Instruments) on a 37-38 °C heating pad. The animal’s eyes were covered with a thick layer of lubricating ointment (Refresh, Alergan Inc.). Using scissors, a flap of skin about 1 cm^2^ in area was retracted from the skull and the gelatinous periosteum was removed with a small scissors. The skull was cleaned and dried with sterile cotton swab. Prior to the implantation, the fiber optic ferrule and electrode pins were disinfected with 70% isopropyl alcohol and allowed to air dry. Coordinates for fiber optic and LFP electrode placement were (AP = −0.2 mm, ML = +1.35 mm, DV = 4.0 mm) (AP = 2.0 mm, ML = 1.0 mm, DV = 0.2 mm) respectively. A custom made headplate was directly affixed with a thin layer of C&B Metabond (Parkell) such that it covered the whole skull and the surrounding area of the ferrules and the electrode pin were fixed with Krazy glue. An electrode was glued on the surface of the skull over the cerebellum as a reference for LFP recording. The skull and the head-plate were secured with a thin layer of Metabond (C&B Metabond) and a layer of dental cement except the stroke area.

**Figure 1.**
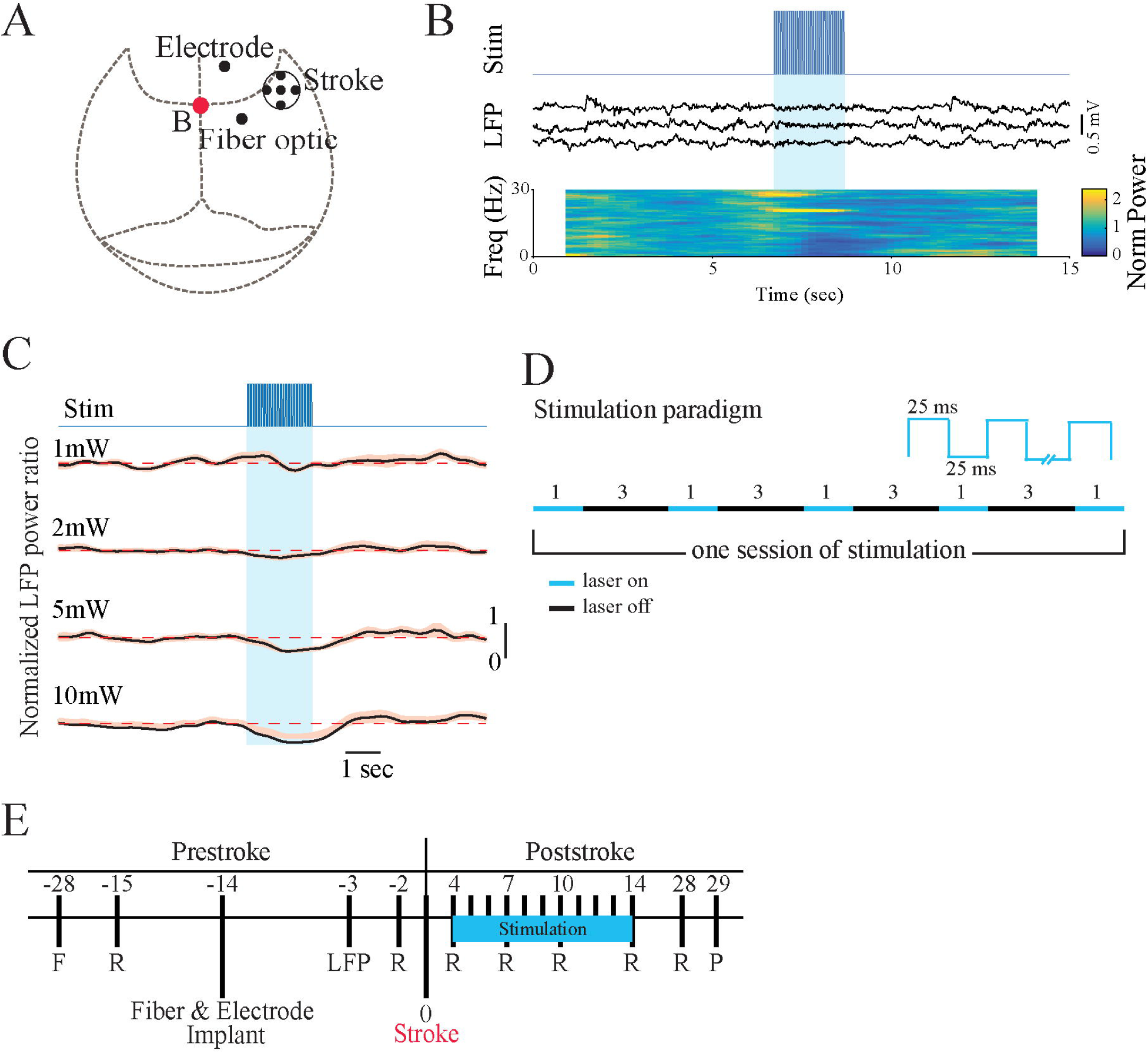
Experimental procedure and stimulation paradigm. (A) Schematic of a mouse skull marked with the location of ischemic stroke, fiber optic and LFP electrode implantation. (B) Stimulation trace shows the onset of laser stimulation. Three examples of LFP recordings from M1 area. Shaded blue area indicates illumination of the nucleus basalis with 25-ms light pulses at 20Hz for 2 sec. Normalized LFP power spectrogram indicates changes in power at each frequency averaged for twenty trials. (C) Difference in the activation level of the cortex following various laser powers measured by mean power ratio from 20 trials. Shaded error bars represent s.e.m. For each plot, y-axis represents normalized power. Dashed lines indicate the baseline prior to stimulation. (D) Stimulation Paradigm. Animals in Stimulation group received five 1-minute laser stimulations with 3-minutes of rest in between. Stimulation was a square pulse at 20 Hz with 25ms on and 25ms off. (E) Experimental timeline before and after stroke induction. All mice were filmed during single pellet reaching task 2 days before stroke and on days 4, 7, 10, 14, and 28 poststroke. Optogenetic stimulation occurred daily from day 4 to day 14 poststroke and after behavioural assessments. F, food restriction; R, reaching task; LFP, local field potential recording; P, perfusion.

### Optogenetic stimulation and LFP recording

The fiber optics were prepared in house following guidance from ThorLabs Manual (FN96A). Briefly, fiber optic cores (Part # FT400UMT, Thor Labs) were cut to 16 mm lengths, inserted into ceramic ferrules (Part # CF440-10, Thor Labs) and cemented in place with epoxy (Part # T120-023-C2, Thor Labs). After allowing the epoxy to set for 24 hours, the “bare” end was cut to 5.0 mm in length with a diamond cutter (Part # S90R, Thor Labs). This “bare” end eventually implanted into the brain. The other end of the fiber optic core was trimmed and polished until flush with the ceramic ferrule and showed an appearance of “polished glass” under a dissecting microscope at 10x power. Before implantation, the ferrules were tested for light transmittance with an optical power meter (Part # PM100D, Thor Labs) by attaching the polished end to a 473 nm laser (Shanghai Dreams Lasers Technology, SDL-473-100T). All fiber optic implanted in the nucleus basalis had an average measured output of ∼5.5mW. Teflon coated 50 µm stainless steel wires (A-M Systems) were used for neocortical LFP recordings. The tip of the electrode was placed in the superficial layers of motor cortex. The local field potential signal was amplified (x1000) and filtered (0.1-10 KHz) using a Grass A.C. pre-amplifier Model P511 (Artisan Technology Group ®, IL) and digitized using a Digidata 1440 (Molecular Device Inc., CA) data acquisition system at 2 KHz sampling rate.

### Analysis of LFPs

Analysis of LFP traces was performed using custom-written code in MATLAB (Mathworks, Natick, MA). LFP signals were down-sampled to 2 kHz, and frequency components of the traces were then extracted by multitaper spectral analysis using the Chronux toolbox ^36^ (http://chronux.org). To measure cortical desynchronization, as a measure of ACh release, the frequency components in 1.5 s time windows that overlapped by 0.1 s were calculated. Normalized power in the 0.1-6 Hz band was calculated by dividing power by the mean 0.1-6 Hz power before the stimulus. Mice were included if desynchronization was observed by optogenetic stimulation of the nucleus basalis cholinergic neurons. For each animal we determined whether there was desynchronization by comparing, for 20 trials, the mean 0.1-6-Hz power 0.5–2.6 s after stimulus onset with the mean power 2 s before the stimulus, using a paired t-test with a P < 0.05 criterion. Effective stimulation was defined by desynchronization of cortical activity as previously described ^37^. On the basis of these experiments (Figure 1B-D), optogenetic stimulation (5 mW, 20 Hz with 25ms on and 25ms off) consisting of five stimulations was selected for the experiments. A series of 1-minute laser stimulation on, at 3-minutes off was given daily from day 4 to day 14 poststroke 3-hr before behavioural testing (Figure 1D).

### Photothrombotic stroke

To induce a forelimb sensory cortex stroke, animals were anesthetized with isoflurane (1-2%), placed on a 37-38 °C heating pad, and the animal’s head-plate was secured with plastic forks on a custom-made surgical plate. The animal’s eyes were covered with a thick layer of lubricating ointment (Refresh, Alergan Inc.). To facilitate photoactivation, a round circle with 1.0mm in diameter on the skull was thinned to ∼ 50% of its original thickness. The center of the circular region was AP = 0.5 mm and ML = 2.5 mm from the Bregma. Photothrombotic stroke was induced by injecting 10 mg/ml of Rose Bengal intraperitoneally 5 minutes prior to 20 minutes of green laser illumination (532 nm wavelength at power 10 mW) at the thinned region as previously described ^38-40^. The thinned area was covered with dental cement. The control mice went through the same procedure except that they received intraperitoneal saline injection rather than Rose Bengal. After the surgery, mice were kept in the recovery room for two days before being returned to their home cage.

### Video recording

Reach behavior were filmed from a frontal view with a Panasonic HDC-SDT750 camera at 60 frames per second at an exposure rate of 1ms. Illumination for filming was obtained by using a two-arm cold light source (Nicon Inc.), with the arms positioned to illuminate the reaching target area from a frontolateral location on each side of the reaching apparatus. Video was replayed frame-by-frame on a personal computer using QuickTime player (10.5 © 2009-2018, Apple Inc) for scoring.

### Reaching apparatus and training and assessment

The reaching box was made of clear plexiglass (19.5-cm long, 8-cm wide, and 20-cm high) with a slit (1-cm wide) located in the center of the front wall ^41, 42^. On the outside of the front wall, a shelf 3.8 cm wide was mounted 1 cm above the floor. Two divots were located on the shelf, one centered on the edge of each side of the slit at a distance of 1 cm from the slit. Food items that the mice reached for were 10-mg food pellets (Catalogue # 1811213, TestDiet) placed singly in a divot. At this location, it is difficult for the mice to obtain food with their tongue, but food can readily be obtained with the contralateral hand because mice pronate their hands with a lateral-to-medial movement that brings the palmar surface of the hand over the divot.

Food restricted animals, maintained at 85% body weight by once daily feeding, were habituated to the reaching apparatus by placing them in the box for 10-min/day for ten days. Pellets were initially available on the reaching box floor and within tongue distance on the shelf. Pellets were gradually removed from the floor and placed further away on the shelf until the mice were forced to use their hands to retrieve the food pellet. The pellets were placed in the right divot, so that the mice were required to reach with the left hand to obtain them. Once reaching, mice received twenty pellets in each training/testing session and each pellet presentation defined a trial. Training was considered complete once each mouse’s success rates reached asymptotic level on three consecutive days of training. Mice were habituated to the laser cable in their home cages during the last 5 days of training.

The reach behavior of mice was subjected to analysis of three measures: success score, movement components, and sensorimotor integration.

#### (1) Success

Three measures of success were used on each trial: (A) Success was defined as a reach in which the mouse successfully grasped the food and brought the food to the mouth for eating (Supplemental Video 1). (B) First trial success was the percent of trials on which a mouse successfully grasped the food on the first attempt and brought the food to the mouth. (C) Attempts were defined as any reach on which a mouse inserted a paw through the aperture.

#### (2) Movement components

Movement components were scored frame-by-frame from the video using a previously described scoring system ^41-43^. Three scorers (BM, RA, AG), who produced highly correlated ratings (r=>90) from the first three successful reaches from each mouse. Twelve components of a reach; hindfeet, front feet, sniff, lift, elbow in, advance, pronation, grasp, supination I, supination II, release, and replace were rated on a three-point scale. A score of 0 was given to a normal movement, a score of 0.5 was given to a movement that is recognizable but incomplete or misplaced, and a score of 1 was given to unidentified or absent movements (for details see ^41^).

#### (3) Sensorimotor integration

Once control mice grasp a food pellet, they indicate that they are successful by bringing the food to the mouth for eating. Three measures of bringing the food to the mouth was used as a measure of sensorimotor integration; did a grasp occur at the end of the hand advance to the food, did the hand remain closed as it was withdrawn, and was the closed hand then brought to the mouth (Supplemental Video 2). After each failed reach, the movements were scored: if a mouse’s hand was closed, it received a score of “1”; if the hand was open, it received a score of “0” for the grasp and withdraw movements; in addition, if the hand was brought to the mouth a score of “1” was given and if it was not a score of “0” was given.

### Histology

Mice were anesthetized and perfused through the heart with 1x phosphate buffered saline followed by 4% paraformaldehyde in 1x phosphate buffered saline. Brains were removed and post-fixed in 4% paraformaldehyde overnight and then cryoprotected in 30% sucrose in 1x phosphate buffered saline and 0.02% sodium azide. The brains were sectioned at 40μm on Blockface and every third section was mounted on charged Superfrost Plus Micro Slide (VWR), stained with Cresyl violet (Nissl staining), and cover slipped using Permount (Fisher-Scientific). Slides were imaged using 20X objective lens with Nanozoomer (2.0RS, Hamamatsu) and stroke size computed as the area of stroke region for each brain slice. To confirm the expression of light sensitive channels tagged with YFP in the cholinergic nuclei, brain slices were subjected to immunohistochemistry for ChAT. Brain sections were fixed on a charged Superfrost Plus Micro Slide (VWR), washed in Tris-buffered Saline (TBS), and blocked in a solution containing 3% goat serum and 0.3% Triton-X in TBS for 2 hours. The slides were incubated in primary antibody, rabbit anti-ChAT (monoclonal, ab178850, Abcam, 1:5000), and TBS with 0.3% Triton-X at room temperature in a dark humid chamber overnight. After the first incubation, the slides were given three 10-minute washes and then incubated with secondary antibody, anti-rabbit-alexa-594 (IgG [H + L]) and TBS with 0.3% Triton-X for 5 hours. After that, the slides were given two 10-minute washes, air dried, covered with coverslips with Vectashield H-1000 (Vector Laboratory), and imaged using Nanozoomer (2.0RS, Hamamatsu).

### Statistical analysis

All behavioural scores were subjected to Shapiro-Wilk test for normality (Supplemental Table 1) and Kruskal-Wallis H test (Supplemental Table 2) using SPSS (v.26.0.0.0). A *p*-value < 0.05 was considered significant. For the stroke size, the mean area of stroke region for the Stim and No Stim groups was subjected to an independent t-test using SPSS (v.26.0.0.0). A *p*-value of less than 0.05 was considered significant. For local field potential analysis, spectrogram of each trial was calculated on the 3-sec long sliding window using custom written program in MATLAB (MathWorks). Spectrograms were averaged across the trials and divided by the baseline calculated from 6 seconds before the optogenetic stimulation.

## Results

### Optical stimulation of the nucleus basalis cholinergic neurons can activate peri-infarct areas

To examine if repeated cholinergic stimulation could activate plasticity-associated mechanisms and functional recovery, a repeated neuronal stimulation paradigm at poststroke days 4–14 (see experimental timeline, Figure 1E) were implemented. A photothrombotic stroke was used to generate infarct in the forelimb somatosensory cortex (S1). Three groups of ChAT-Cre/Ai32 (ChR2-YFP) transgenic mice, were used: sham stroke + no stim, stroke + no stim, and stroke + stim. Staining for ChAT-immuno responses and its overlap with ChAT-Ai32-EYPF of the region are shown in Figure 2A, indicated high levels of ChR2 in cholinergic neurons. An optical fiber was stereotaxically implanted in the nucleus basalis. The placement of the fiber optic implant in the mice was confirmed by examining the location of the end of the implant in the Nissl stained brain sections in relation to the Paxinos mouse brain atlas ^44^ and showed that the fiber optics were placed in the nucleus basalis (Figure 2B). We first examined if reliable cortical activation could be achieved by optical stimulation paradigm consisting of 5 successive 1-min laser stimulations, separated by 3-min rest intervals (Figure 1 D). In vivo electrophysiological LFP electrode recording from primary motor cortex (M1) in isoflurane-anesthetized mice indicated that this stimulation paradigm could generate reliable and consistent cortical desynchronization in all 5 stimulations. The normalized power spectrogram of LFP signal reveals a decrease in power following the stimulus (Figure 1B; dark blue regions). Higher level of cortical desynchronization was observed when laser intensity increased from 1 to 10 mW (Figure 1C). Desynchronization was not observed in negative ChAT-cre/Ai32 mice (n=3, data not shown). Stroke size and location was confirmed after brain sectioning and staining with Cresyl violet (Figure 2C). The stroke sites consisted of damage to the primary somatosensory forelimb area within cortical layers I to VI, above the external capsule. There was no statistically significant difference in the area of stroke for Stim vs No stim groups (t(15) = 0.582, *p* > 0.05; Figure2D).

**Figure 2.**
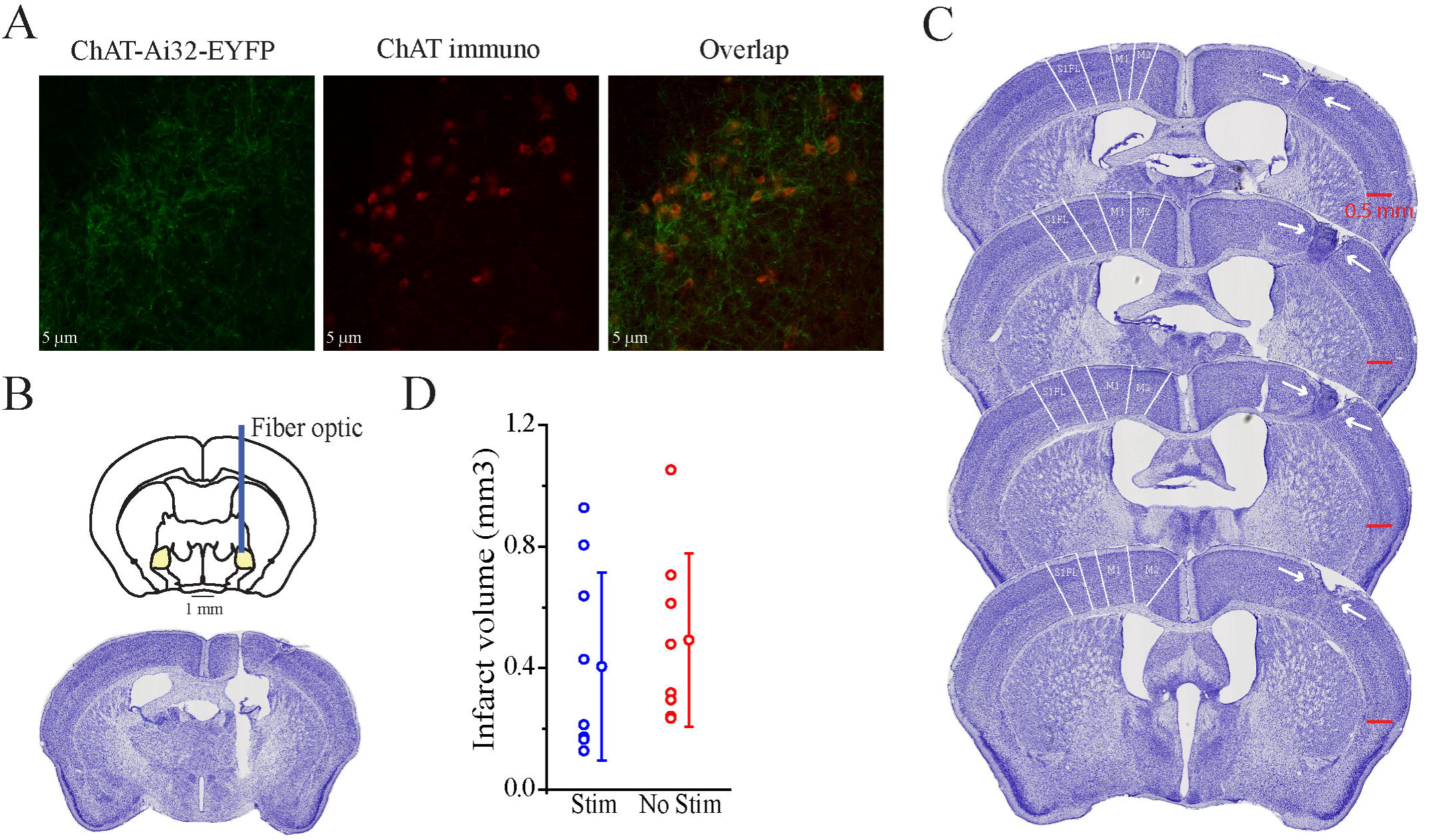
No effect of optogenetic stimulation of the nucleus basalis cholinergic neurons on infarct volume. (A) example of ChR2-EYFP fluorescence (left), ChAT immunoreactivity (middle), and merged image (right) in coronal sections of the nucleus basalis. (B) Fiber optic implant site in the nucleus basalis. (C) Representative montage of cresyl violet stained coronal brain sections showing the anterior to posterior extent of photothrombotic stroke on the right hemisphere. Stroke site includes the primary somatosensory forelimb area. (D) Stroke volume for Stim (n=9) and No Stim (n=8) groups.

### Repeated optogenetic stimulation of NB of cholinergic neurons improves number of successful reach performance after stroke

To address whether repeated ipsilesional cholinergic system stimulations can promote functional recovery, we evaluated the behavioral performance of stroke mice on the single pellet skilled reaching task, a sensitive and reproducible sensory-motor behavior test used to detect neurological deficit after stroke ^45, 46^. Because the single pellet reaching task requires the use of one hand to reach for and grasp a food pellet and bring it to the mouth for consumption ^41^, performance was measured using endpoint scores of the affected hand, ratings of the movement components of reaching and sensorimotor integration assessment. Sensorimotor integration assessment measured successful recognition that there is or, is not, food in the hand after a reach by recording whether a mouse brought its hand to its mouth after a reach. In previous studies of rodents, these measures have provided a definitive measure of poststroke performance ^47-49^. The prestroke assessment of reaching, as assessed with measure of overall Success, First Trial Success, and number of Attempts (See methods), showed that there was no Group difference on any of these measures, (H(2)=2.397, p=0.302; H(2)=2.013, p=0.366; H(2)=5.203, p=0.074; Supplemental Table 2). On the poststroke recovery performance, the Stim group displayed better performance than the No Stim group, and on some measures the Stim group did not differ from the Sham group, as confirmed by the following statistical analyses.

A Kruskal-Wallis H test on poststroke Success performance (Figure 3A) gave a significant effect for poststroke Day 4 and 7 (H(2)=8.816, p=0.012 & H(2)=9.757, p=0.008 respectively; Supplemental Table 2). Follow up pairwise comparison of the experimental groups showed significant difference on poststroke Day 7 (Supplemental Table 3). No Stim group was impaired relative to Sham (p=0.007) and Stim group (p=0.006), but the Sham and Stim groups did not differ (p=0.987). Kruskal-Wallis H test on First Reach Success performance (Figure 3B) gave no significant effect for Days 4, 10, 14, and 28 poststroke (for details see Supplemental Table 2). Kruskal-Wallis H test on total Attempts (Figure 3C) gave no significant effect for Days 4, 7, 10, 14, and 28 poststroke (for details see Supplemental Table 2).

**Figure 3.**
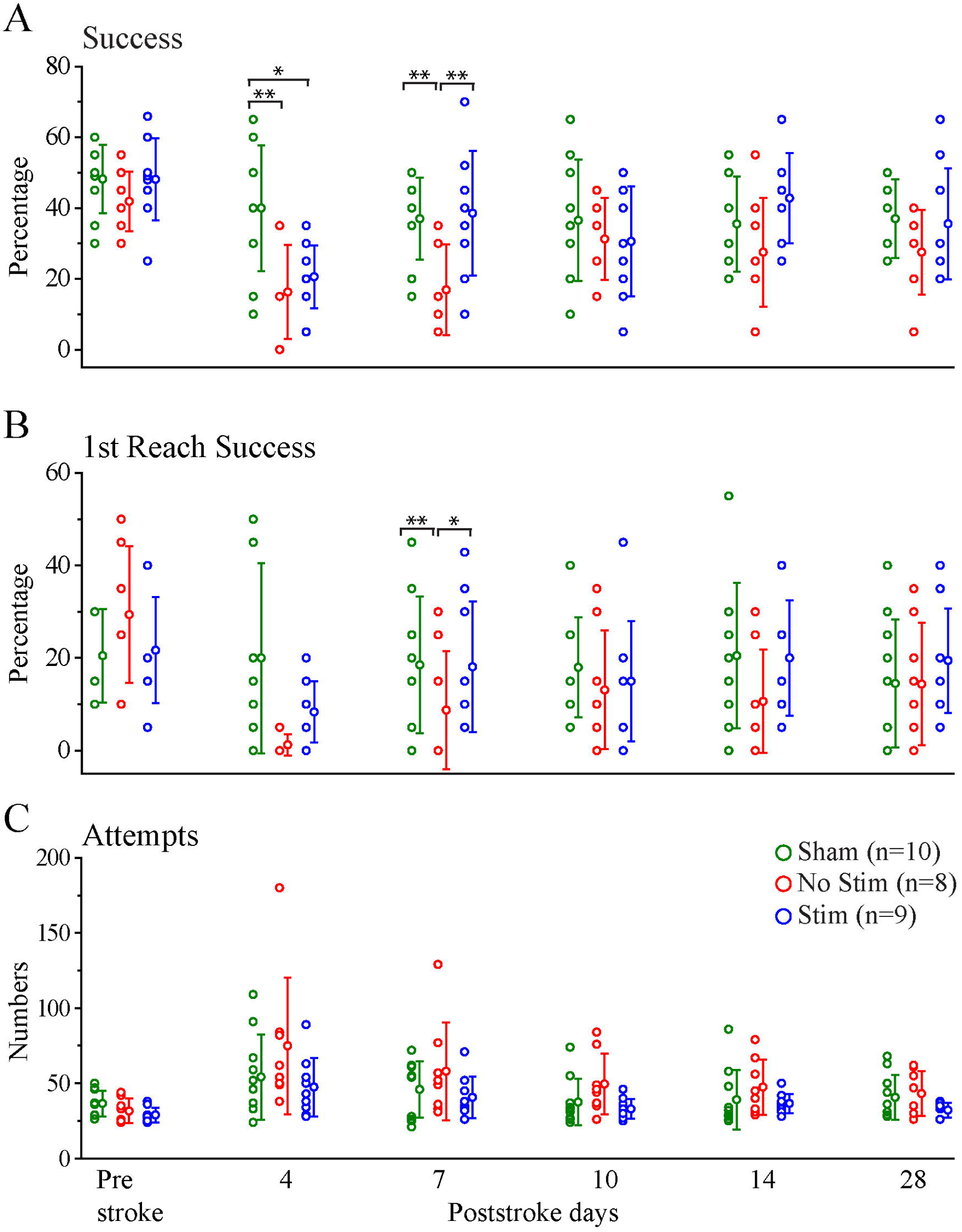
Recovery of endpoint measures is enhanced by optogenetic stimulation. Endpoint measures (data points and SD) for Stim, No Stim, and Sham Groups on prestroke and poststroke days 4, 7, 10, 14, and 28. (A) Success percentage or hit percent. (B) Success percentage on first reach attempt. (C) Attempts. (\**p*<0.05, \*\**p*<0.01, \*\*\**p*<0.0001).

### Optogenetic stimulation of NB cholinergic neurons improves reach quality after stroke

To assess the effect of stroke and NB optogenetic stimulation on the quality of reaches, 12 components of movements were scored on a 0, 0.5, and 1 scale (good, impaired, and absent performance) from the first three success reaches. A higher score indicates inferior performance. The twelve movement components were then divided into three categories; Posture (symmetry of hind and front feet and sniffing to locate the food), Advance (limb lift, elbow in, advance pronation and grasp), and Withdraw (supination I, supination II, food release and hand replace on floor). The analyses on poststroke performance revealed poorer performance of the stroke groups (combined Stim and No Stim experimental groups) relative to the Sham group and improvement in the performance of the stroke groups over days (Supplemental Table 2). The Advance measure indicated that the Stimulation group displayed faster recovery than the No Stim group. For Withdraw, the stimulation group was better than the No Stim group. These results were confirmed by the following statistical analyses.

After stroke, the Posture measure showed (Figure 4A) a significant effect for Days 7 and 28 poststroke (H(2)=8.404, p= 0.015; H(2)=10.427, p=0.005). Follow-up pairwise comparisons of experimental groups showed that the Sham group had higher scores than both the No Stim (p=0.029) and the Stim group (p=0.007) on Day7 poststroke, but on Day 28 the Sham and No Stim groups had higher scores and the Stim group had significantly lower score (for details see Supplemental Table 3). For Advance (Figure 4B), there was a significant effect for poststroke Day 4 and 14 (H(2)=7.852, p=0.02). Follow-up pairwise comparisons of experimental groups on poststroke Day 14 revealed an effect of optogenetic stimulation (significant difference between No Stim and Sham groups (p=0.005), while the Stim and Sham groups show no significant difference (p=0.342); for details see Supplemental Table 3). On the Withdraw measure (Figure 4C), there was a significant effect for poststroke Day 4, 10 and 28 (H(2)=9.345, p=0.009; H(2)=7.817, p=0.02; H(2)=6.847, p=0.033 respectively). Follow-up tests indicated that the Sham and Stimulation groups did not differ, and their scores were lower than the No Stim group (for details see Supplemental Table 3). Overall, these results suggest a positive impact of Nucleus Basalis optogenetic stimulation of cholinergic neurons on quality of reach performance after stroke.

**Figure 4.**
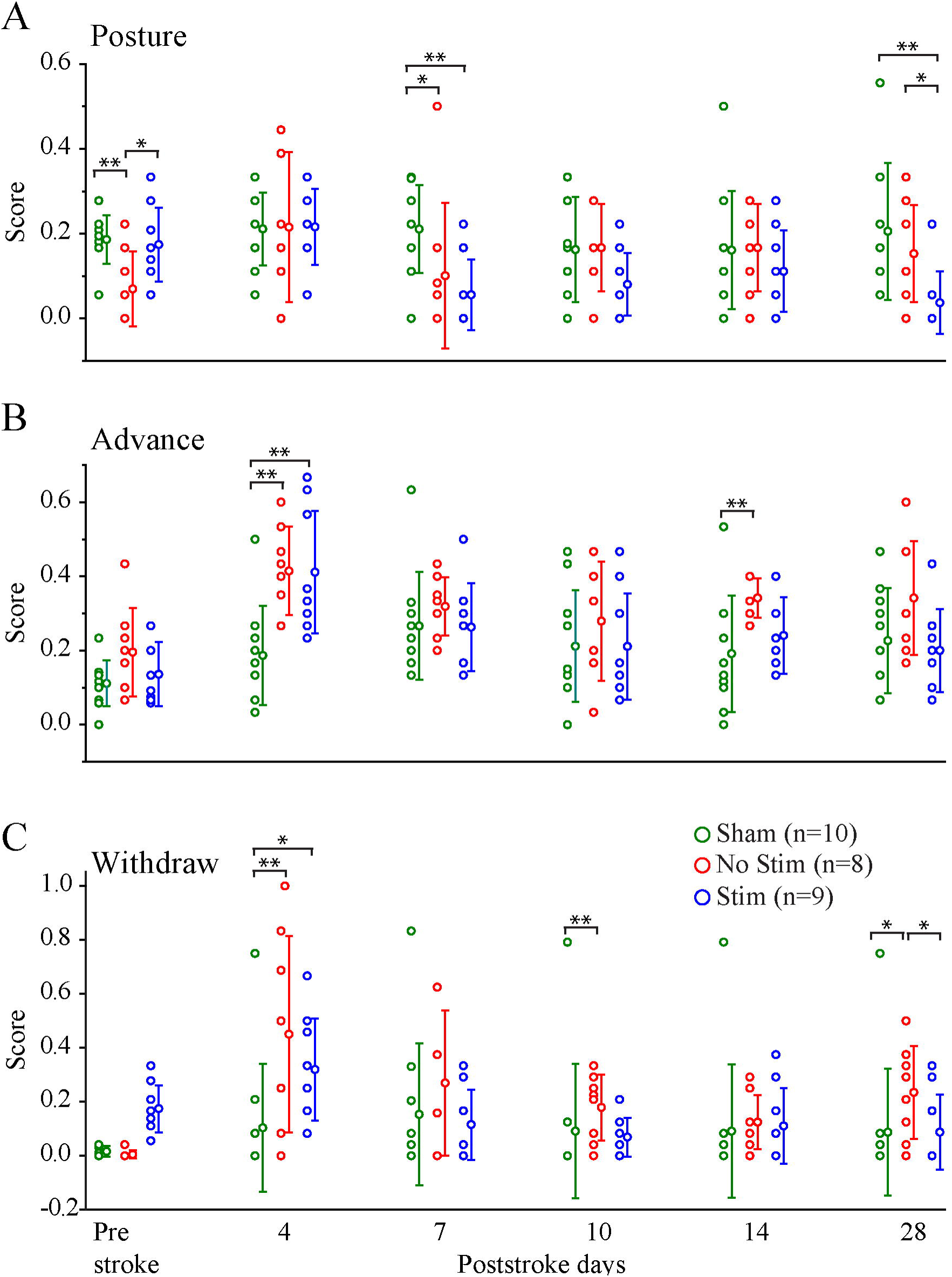
Quality of skilled reaching behaviour is improved by optogenetic stimulation of NB cholinergic neurons. Twelve movement components categorized as posture, advance, and withdraw (data points and SD) for Stim, No Stim, and Sham Stroke Groups on prestroke and poststroke days 4, 7, 10, 14, and 28. Scores are in a scale of 0 to 1 where 0 indicates normal movement and 1 indicates impairment. (\**p*<0.05, \*\**p*<0.01, \*\*\**p*<0.0001).

### Optogenetic stimulation of NB cholinergic neurons does not improve recovery of sensorimotor integration after stroke

Before stroke, there were no differences on measures of Grasp, Retract Close and Hand to Mouth among the experimental Groups (H(2)=2.970, p=0.226; H(2)=2.806, p=0.246; H(2)=2.344, p=0.310). On successful reaches, mice grasped, retracted, and brought their hand to their mouth (Supplemental Video 1), while on unsuccessful reaches they did not do so (Figure 5A; Supplemental Video 2). After stroke, mice in the stroke groups were more likely to close their hands after a missed grasp, withdraw the closed hand and make a movement of bringing the hand to the mouth as if to take food from it (Figure 5B; Supplemental Video 2), a behavior termed fictive eating ^50^. Supplemental Video 2 illustrates the typical absence of grasping for a Sham mouse and the presence of fictive eating in a mouse with sensory forelimb stroke. Using Grasp, Retract Close and Hand to Mouth measurement, we found that the impairment was very similar in the No Stim and the Stim stroke groups. After stroke for Grasp, Retract Close, and Hand to Mouth measures (Figure 6A-C), there were significant effects for Days 4, 7, 10, 14, and 28 with pairwise comparisons of experimental groups revealing that the stroke groups (both Stim and No Stim groups) had more grasps, retract close, and hand to mouth (p<0.05) (for details see Supplemental Tables 2 and 3) than the Sham group. These results confirm that that optogenetic stimulation of NB cholinergic neurons does not improve recovery of sensorimotor integration after stroke.

**Figure 5.**
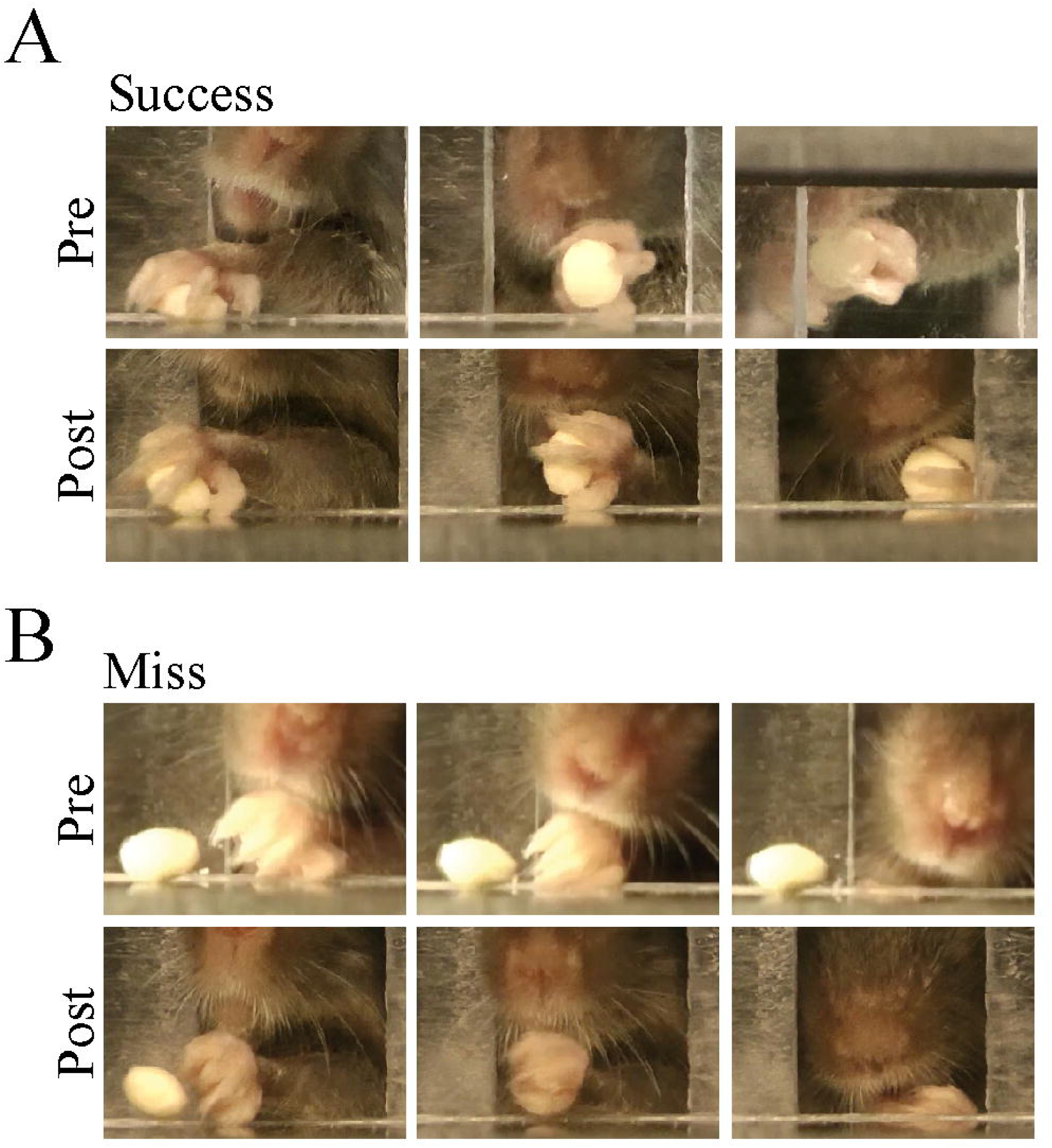
Grasp, withdraw, and hand to mouth during (A) success and (B) miss pre and poststroke. In miss condition prestroke after reaching, the mouse does not close digits (left) and withdraws the hand without supinating (middle). Once the hand gets to the end of the opening, the mouse does not check the hand for food (right.). In miss condition poststroke after reaching, the mouse closes digits (left) and withdraws the hand while supinating (middle). Once the hand gets to the end of opening, the mouse sniffs the hand and checks it for food (right).

**Figure 6.**
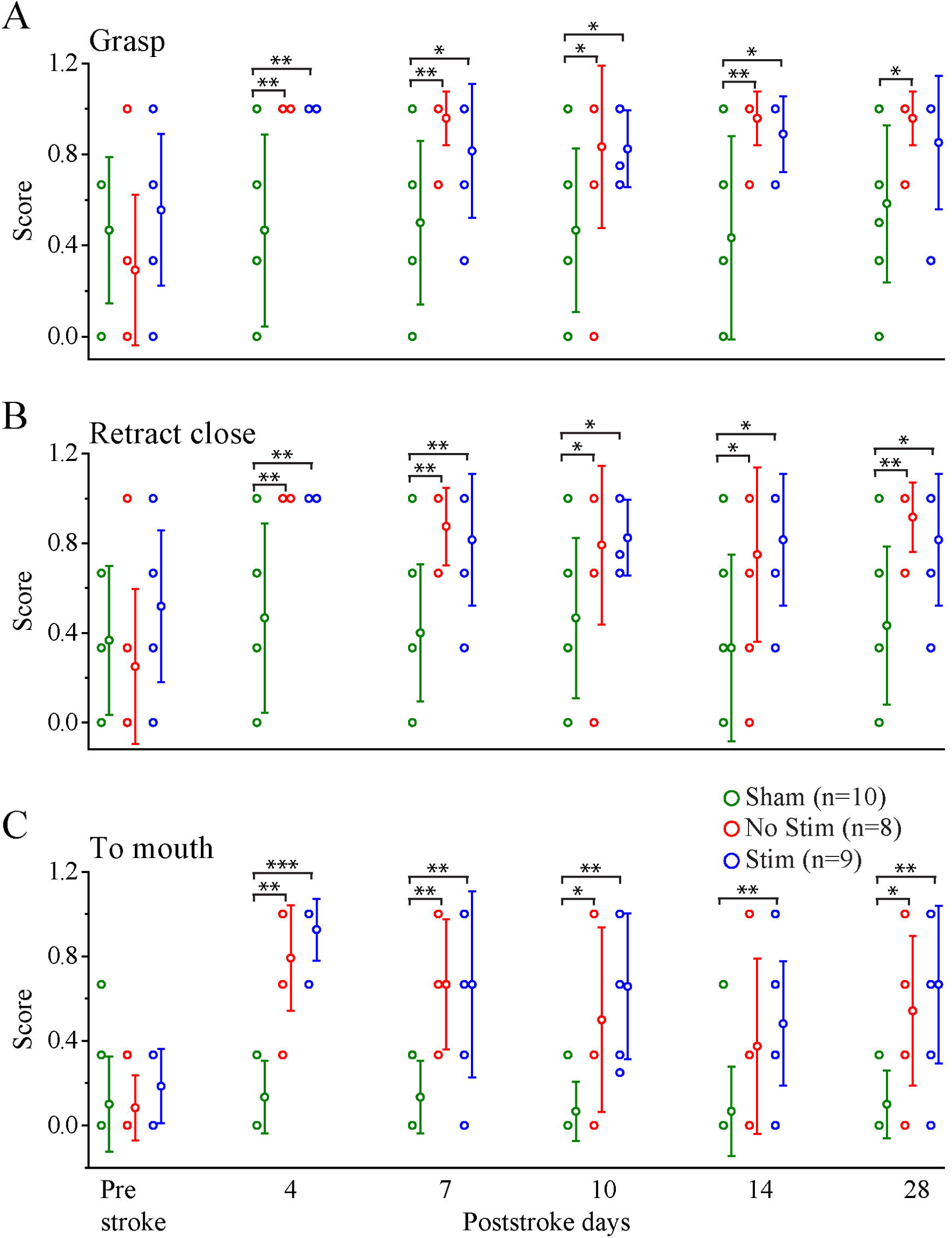
Stroke-impaired sensorimotor integration is not improved by optogenetic stimulation of NB cholinergic neurons. Occurrence of three behaviours Grasp, Retract Close, and Hand to Mouth (data points and SD) for Stim, No Stim, and Sham Groups on prestroke and poststroke days 4, 7, 10, 14, and 28 in Miss condition. (\**p*<0.05, \*\**p*<0.01, \*\*\**p*<0.0001).

## Discussion

There is abundant evidence that the basal forebrain cholinergic system, as an essential component of the neuromodulatory system, plays an important role in cellular excitability, synaptic plasticity, improvement of sensory coding, control of behavior state and cognitive function. Cholinergic neurons project their axons to many areas of the brain and act on a variety of receptors and target cells, therefore modulation of ACh receptors result in complex and often opposing influences on numerous biological processes including poststroke synaptic plasticity. In this study, using longitudinal repetitive optogenetic stimulation, in vivo electrophysiology, and animal behavior, we explored the role of cholinergic circuits in functional plasticity and behavioral recovery days and weeks after stroke.

Selective photothrombotic stroke to the forelimb sensory region of the mouse neocortex produced transient impairments in end point measures and reach movements and a chronic impairment in sensory integration in mouse performance on the single pellet reaching task. Optogenetic stimulation of the nucleus basalis cholinergic systems during the poststroke period improved some measures of recovery on end point measures and movement but was without effect on measures of sensory integration. These results suggest that the upregulation of cholinergic tone can facilitate some aspects of behavioral recovery and confirm that cholinergic projections to the sensorimotor cortex play a role in behavioral plasticity.

The design of the present study was based on Cheng et al ^30^, who report that optogenetic stimulation of ipsilesional motor cortex improved recovery from middle cerebral artery stroke on end point scores of a test of walking on a rotating rod. In the Cheng et al ^30^ study, stimulation increased cerebral blood flow, the neurovascular coupling response, and expression of activity-dependent neurotrophins, including brain-derived neurotrophic factor, nerve growth factor, and neurotrophin. Thus, in the present experiment, similar dose and period of optogenetic stimulation were given to mice during the recovery period, except mice underwent selective stroke of the forelimb region of somatosensory cortex and stimulation was delivered to the cholinergic neurons in the nucleus basalis. The nucleus basalis contains cholinergic neurons that selectively project to the mouse sensorimotor cortex and this projection has been proposed to play a role in cortical plasticity and recovery of function after stroke. Prior to stimulation, the effect of optogenetic stimulation was assessed for its effects on cortical LFP signal desynchronization ^37, 51^. In addition, the mice were tested in a more comprehensive behavioural test, a single pellet reaching task that evaluates end point measures as well as motor and sensory function.

Evidence from anatomical and electrophysiological studies of primates show that somatosensory and motor cortex have a shared role in the control of skilled movement. Electrical stimulation of both regions produces reach and grasp movement and there are rich anatomical interconnections between sensory and motor cortical regions ^52, 53^. There have been no similar studies on mouse sensorimotor cortex ^54^ but the present results show that selective stroke of the forelimb area of sensory cortex results in some of the same deficits as those described following selective stroke to the mouse forelimb region of motor cortex ^41^. The present sensory forelimb stroke transiently impaired end point measures on a reach-to-eat task and produces impairments in many of the movements of reaching, just as does forelimb stroke to motor cortex. The mice with forelimb sensory stroke also displayed an impairment in sensorimotor integration in that they were just as likely to bring their hand to their mouth, as if to transfer a grasped food pellet, on failed reach attempts as on successful reach attempts. A similar impairment in sensorimotor integration is not reported after motor cortex forelimb stroke in mice but has been previously reported after forelimb sensory cortex stroke in the rat ^55, 56^. Thus, consistent with findings from nonhuman primate and human studies of sensory cortex stroke, forelimb sensory cortex of the mouse contributes to both motor and sensory control of the forelimb in mice. The chronic impairment in sensorimotor integration is not surprising as previous research indicates that sensory function is unlikely to recover after somatosensory stroke ^57-59^. Thus, it can be concluded that the optogenetic stimulation of cholinergic system was effective because it enhanced function in the motor cortex, the likely substrate of movement success and movement quality, which was left intact by the somatosensory forelimb stroke.

The optogenetic stimulation of the nucleus basalis cholinergic neurons was associated with enhanced recovery of reach success and attempts and some of the measures of motor performance, especially advancing the limb to grasp food, but had no effect on measures of sensorimotor integration. The finding supports the idea that cholinergic innervation of sensorimotor cortex is involved in cortical functional recovery and may be beneficial on the recovery of some aspects of behavior. There are two potential explanation for the effectiveness of optogenetic stimulation in improving end point measures of reach behavior. First, following stroke induction in the forelimb somatosensory area, the cholinergic terminals in the peri-infarct region of the cortex may be damaged/depressed resulting in reduction of ACh function in those regions of the cortex that mediated recovery of behavior as it was shown previously ^29^. Stimulation of the nucleus basalis may induce the remaining cholinergic terminals in the cortex release more ACh, thus facilitating plasticity in the motor cortex and peri-infarct region. Second, after stroke induction in the forelimb somatosensory area, the cholinergic neurons in the nucleus basalis that are remote from the stroke site may undergo diaschisis ^60^, resulting in depression of ACh release from the axonal terminals in the cortex. In this respect it is interesting that a number of early studies reporting a transient depression in cholinergic related EEG activity after forebrain and brainstem lesions ^61, 62^. Optogenetic stimulation of the nucleus basalis may boost the level of released acetylcholine by cholinergic neurons, thus dissipating diaschisis.

There are caveats related to the present application of optogenetic stimulation relative to its effectiveness in improving recovery from stroke. First, stimulation was given before and separate from the behavioral assessments as the presence of stimulation during behavioural assessments could interfere with animal performance. The optimal time to begin rehabilitation interventions poststroke is still not well established, and the behavioral changes in skilled reaching following stroke are complex ^63^. Accordingly, there are many potential paradigms for nucleus basalis stimulation delivery that could be explored, including whether stimulation is best if given before, during or immediately after the behavioural assessment tests. Second, cortical desynchronization was used to assess optogenetic stimulation effectiveness as based on previous work ^37, 51^, however, that is only an indirect measure of the ACh release and future work could make more direct measures ^64^. Importantly, however, the results of the present study did show that the stimulation was not harmful because on all behavioral measures the mice receiving stimulation were not impaired relative to the No Stim group. Third, some evidence suggests that therapy within 24-48 hours of stroke in the mouse can be harmful ^65^ and so here optogenetic stimulation and behavioral recovery assessment began on Day 4 poststroke and the maximal benefit of optogenetic stimulation was obtained on Day 7 poststroke on the measure of reach success. This present finding suggests that in future work it would be worthwhile to initiate optogenetic stimulation of cholinergic neurons and behavioral assessment as early as the first day after stroke^66^.

In conclusion, optical stimulation of the nucleus basalis cholinergic neurons following focal ischemic stroke to the primary forelimb somatosensory area restored some but not all features of skilled reaching. There is evidence that some aspects of behavior depend upon the integrity of cortical circuits and although animals may use compensatory movements, the recovery of normal movement and sensory behavior will depend upon proper replacement of lost connections. Nevertheless, our finding here suggests one avenue for improving poststroke functional recovery.

## Author contributions

BM, IQW, and MHM designed the study. BM, and RA performed the experimental methods, trained the animals, and collected the data. IQW, BM, RA, MN, and AG analyzed the data. BM, IQW, and MHM wrote the manuscript to which all authors edited. MHM acquired the funding and provided resources. MHM and IQW supervised the study.

## Acknowledgement

We thank Di Shao for animal husbandry and Jamshid Faraji for help in initial experiments. This work was supported by Canadian Institute for Health Research (CIHR) Graduate Scholarship Program to BM, Natural Sciences and Engineering Research Council of Canada Undergraduate Scholarship Program to RA, Alberta Innovate for Health Science Undergraduate Scholorship Program to AG, Canadian Institutes of Health Research (CIHR) Grant# 390930, Natural Sciences and Engineering Research Council of Canada (NSERC) Discovery Grant #40352, Alberta Innovates (CAIP Chair) Grant #43568, Alberta Alzheimer Research Program Grant # PAZ15010 and PAZ17010, and Alzheimer Society of Canada Grant# 43674 to MHM.

