## Supplemental Tables for "Cholinergic upregulation by optogenetic stimulation of nucleus basalis after photothrombotic stroke in forelimb somatosensory cortex improves endpoint and motor but not sensory control of skilled reaching in mice"

**Table 1**. Shapiro-Wilk test of normality. (note: p<0.05 indicates that data is not normally distributed)
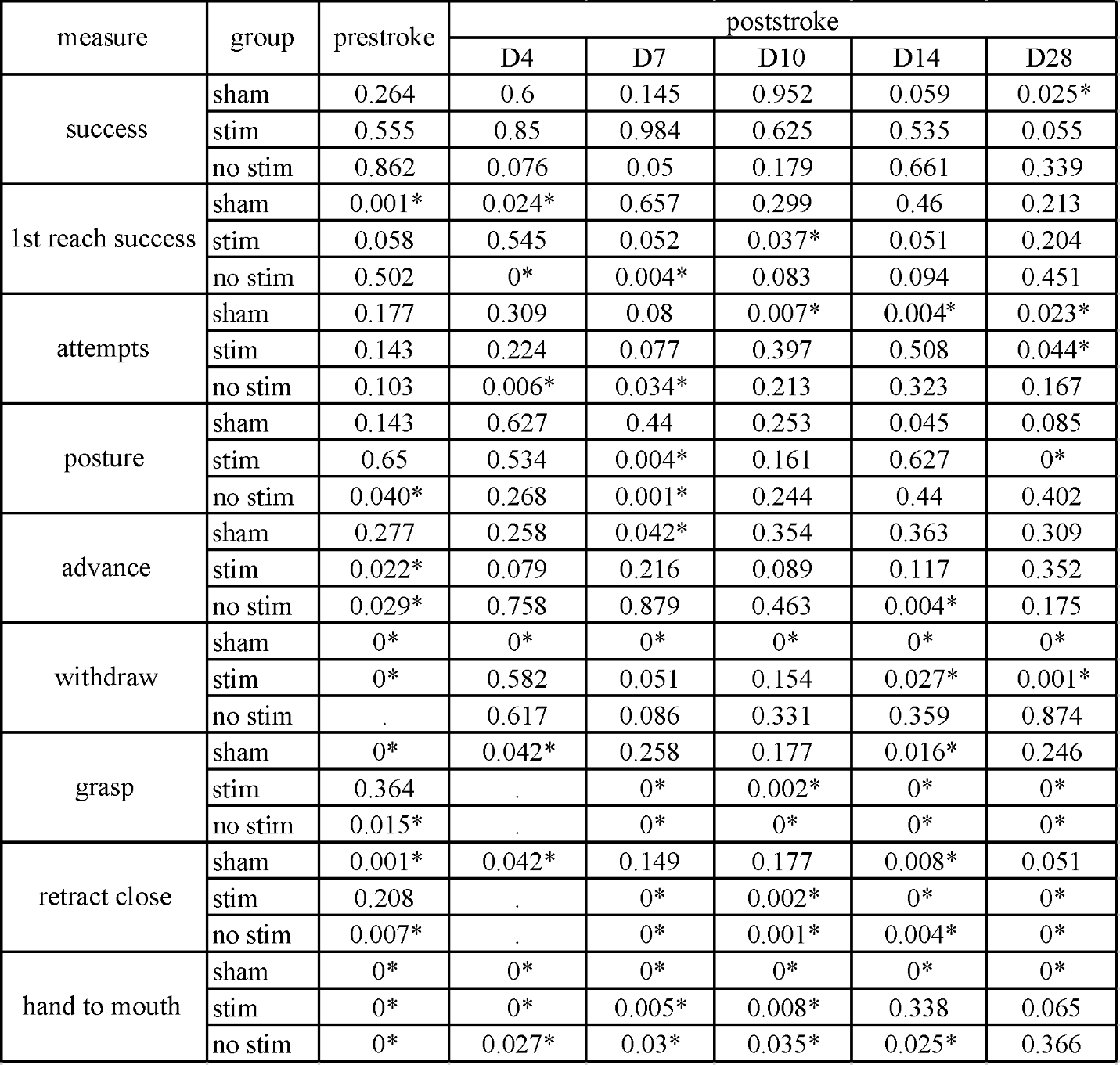

**Table 2**. Kruskal-Wallis H test results.

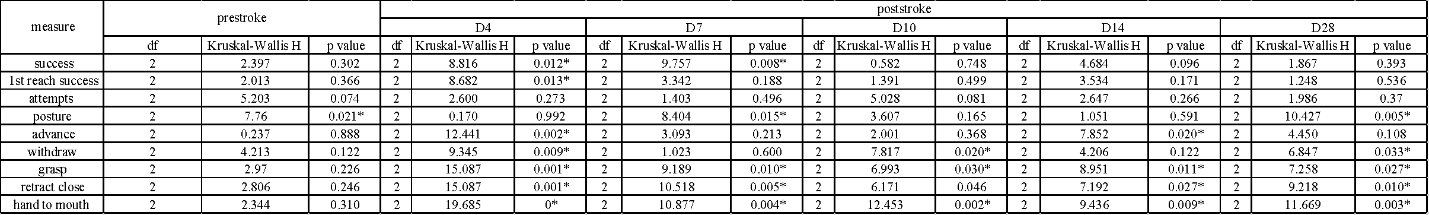

**Table 3**. Pairwise comparison of groups.

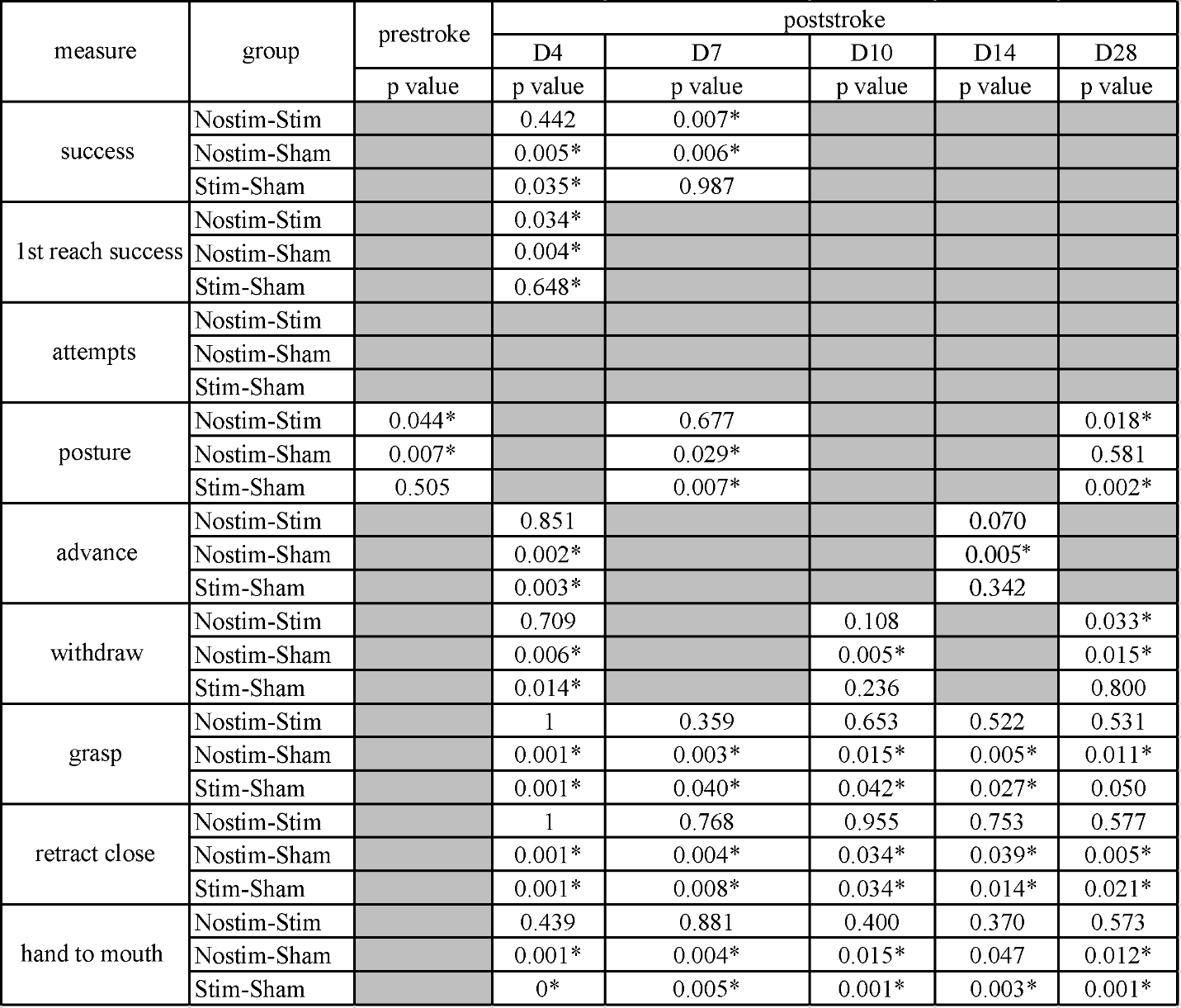

**Supplemental Video Captions**

**Video 1**. Successful reach of a mouse in prestroke and poststroke day 4. The speed of video play is slowed down to 10% of its original speed.

**Video 2**. Attempt in a miss condition prestroke and poststroke day 4 highlighting three measures of sensorimotor integration: grasp, retract close, and hand to mouth. The speed of video play is slowed down to 10% of its original speed.
